# Learning phenotype associated signature in spatial transcriptomics with PASSAGE

**DOI:** 10.1101/2024.09.06.611564

**Authors:** Chen-Kai Guo, Chen-Rui Xia, Guangdun Peng, Zhi-Jie Cao, Ge Gao

## Abstract

Spatially resolved transcriptomics (SRT) is poised to advance our understanding of cellular organization within complex tissues under various physiological and pathological conditions at unprecedented resolution. Despite the development of numerous computational tools that facilitate the automatic identification of statistically significant intra-/inter-slice patterns (like spatial domains), these methods typically operate in an unsupervised manner, without leveraging sample characteristics like physiological/pathological states. Here we present **PASSAGE** (**P**henotype **A**ssociated **S**patial **S**ignature **A**nalysis with **G**raph-based **E**mbedding), a rationally-designed deep learning framework for characterizing phenotype-associated signatures across multiple heterogeneous spatial slices effectively. In addition to its outstanding performance in systematic benchmarks, we have demonstrated PASSAGE’s unique capability in identifying sophisticated signatures in multiple real-world datasets. The full package of PASSAGE is available at https://github.com/gao-lab/PASSAGE.

## Introduction

Spatially resolved transcriptomic (SRT) technologies allow for the profiling of genes expression within their native spatial contexts across a wide range of tissue types^[1–8]^. Existing computational methods for analyzing spatial transcriptomics data have primarily focused on the unsupervised exploration of spatial patterns, such as identifying spatially variable genes, defining spatial domains^[9–12]^, and aligning multiple SRT slices^[11,13–15]^. Meanwhile, benefitting from technical advances and broader application, SRT data are now being generated at increasing volumes across diverse conditions, including both physiological and pathological tissues^[7,16]^, enabling a unique opportunity to systematically identifying signatures associated with specific phenotypic characteristics.

Here, we introduce **PASSAGE** (**P**henotype **A**ssociated **S**patial **S**ignature **A**nalysis with **G**raph-based **E**mbedding), a novel supervised representation learning model designed for large cohorts of phenotypically labeled spatial transcriptomics slices. Combining graph attention auto-encoder (GATE)-based cell/spot-level spatial encoding with slice-level information aggregation through a dedicated attention pooling strategy, PASSAGE achieves accurate classification and clustering of heterogeneous slices, and effectively pinpoints phenotype-associated signatures across multiple slices. PASSAGE is publicly accessible at https://github.com/gao-lab/PASSAGE.

## Results

To effectively utilize spatial information, PASSAGE begins by modeling each slice as a spatial neighbor graph, and employs a graph attention auto-encoder (GATE) to learn spatially aware spot/cell embeddings within each slice. These embeddings capture information not only from the expression profile of individual cells/spots, but also from their spatial neighborhood within a tissue context. PASSAGE then introduces a dedicated attention pooling layer that aggregates the embeddings of all cells/spots within each slice into a single slice-level embedding (**Methods**), which functions as a learnable dynamic averaging process capable of focusing on specific spatial regions. The attention pooling part could be further trained using triplet-based contrastive learning, supervised by phenotypic annotation of the slices, to “guide” effective attention to regions most relevant to phenotypic differences (**Figure 1**).

**Figure 1.**
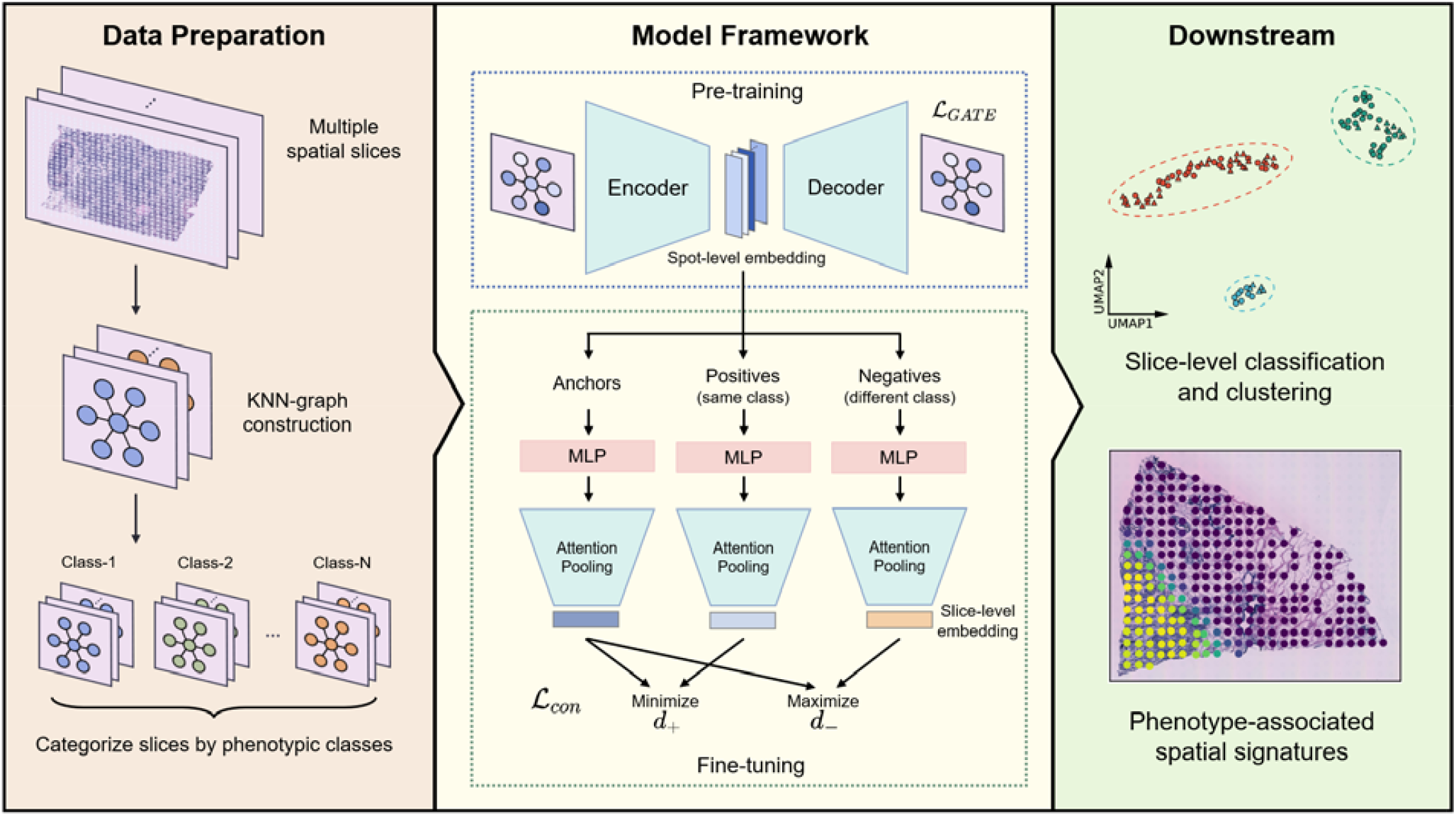
Overview of PASSAGE. The workflow of PASSAGE comprises three main steps: data preparation, model training, and downstream tasks. In data preparation, PASSAGE constructs a spatial neighbor graph for each spatial slice. Then, the GATE framework is employed to learn spot-level embeddings. Finally, guided by phenotype-supervised contrastive learning, the spot-level embeddings are dynamically aggregated using a learnable attention pooling layer to obtain slice-level embeddings that best distinguishes phenotypic labels. The trained model can subsequently be utilized for slice classification, clustering, and identification of phenotype-associated spatial signatures.

### Systematic benchmarking shows outstanding performance of PASSAGE

To conduct benchmarks with definitive ground truth, we generated two synthetic spatial datasets with varying levels of complexity. The first dataset (synthetic data 1) comprised slices of two phenotypic classes, used as the supervision label for the compared algorithms. Each class contains two simulated cell types. The two classes were distinguished by whether these two cell types are spatially separated (class 1) or infiltrated each other (class 2) (**Figure 2A**). The cell types exhibit mild discernability in their expression profiles, mirroring the characteristics of current *in situ* spatial transcriptomics technologies, where the transcriptomic differences captured between cell types are less pronounced than those in conventional single-cell data due to limited spatial resolution or segmentation inaccuracies (**Figure S1**).

**Figure 2.**
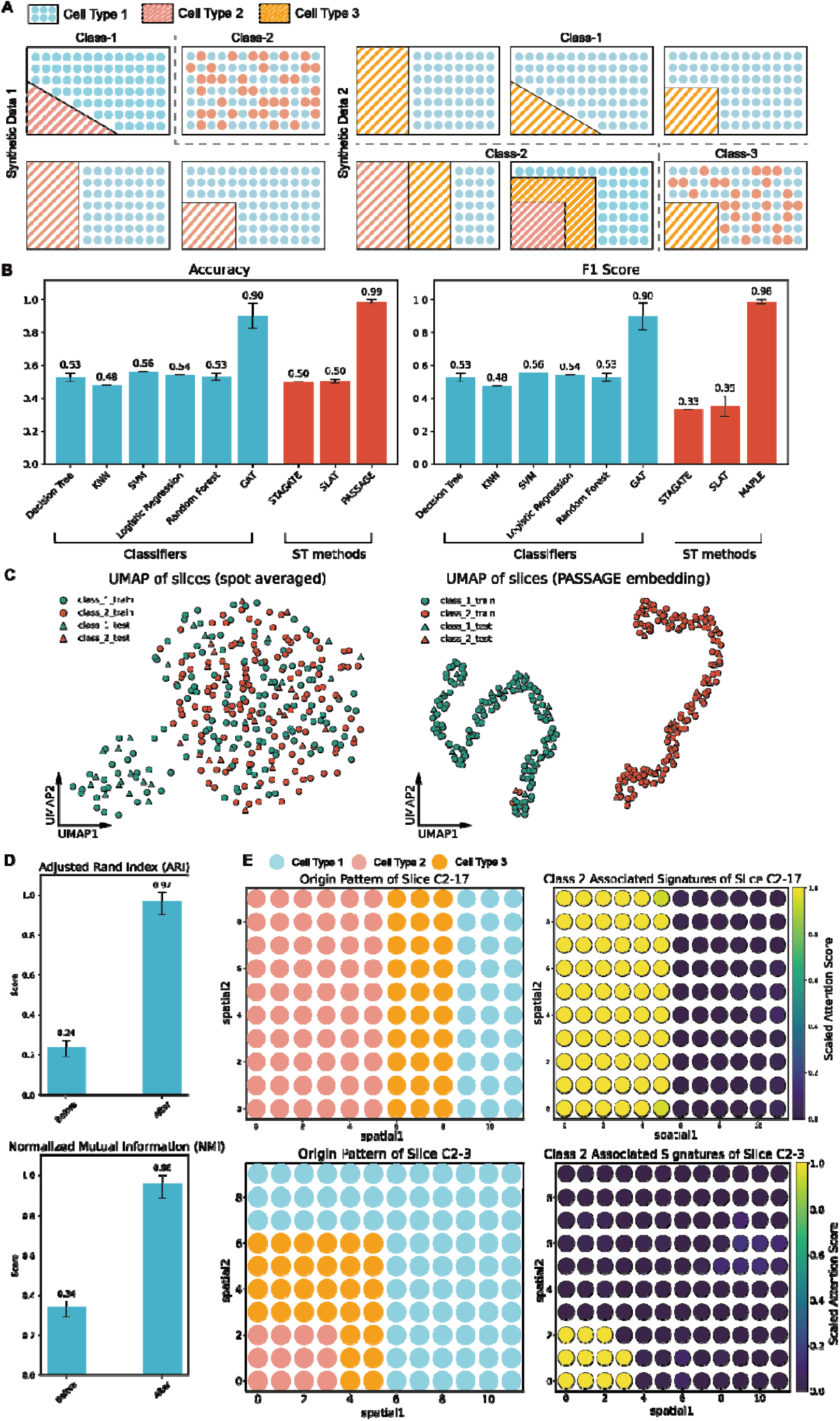
Systematic simulation study. A) Cell type spatial patterns in synthetic data 1 (left) and 2 (right). B) Comparison of classification metrics for PASSAGE and other baseline models in synthetic data 1. C) UMAP visualization of slice embeddings obtained by simply averaging all spots (left) or using PASSAGE (right). D) Clustering metrics for averaging-based embeddings (top) and PASSAGE embeddings (bottom) in synthetic data E) Examples of class-2 associated spatial signatures identified by PASSAGE in synthetic data 2. The error bars indicate mean ± s.d.

To the best of our knowledge, PASSAGE is the first method specifically designed based on supervised classification of spatial transcriptomics slices, so there are no directly comparable methods for benchmarking. Thus, we evaluated PASSAGE against two categories of relevant methods. The first category includes general-purpose classification algorithms, including decision trees, k-nearest neighbors (k-NN), support vector machines (SVM), logistic regression, random forests and graph attention networks (GAT).^[10,11,13]^ The second category consists of unsupervised representation learning algorithms tailored for spatial transcriptomics data, including STAGATE and SLAT (**Methods**).

PASSAGE achieved substantially higher classification accuracy compared to the other methods (**Figure 2B, S2**). Even though the two classes of synthetic slices were intermingled in their original UMAP embedding space obtained by averaging all spots in each slice (**Figure 2C**), slice-level PASSAGE embeddings clustered neatly into two well-separated categories that closely match the true class labels (**Figure 2C**), as also reflected by a significant increase in ARI (Adjusted Rand Index) and NMI (Normalized Mutual Information) (**Figure 2D**). These results indicate that PASSAGE effectively distinguishes spatial slices belonging to different phenotypic groups within its embedding space. Of note, PASSAGE is designed to learn slice-wise embeddings (rather than spot/cell-wise ones of SLAT and STAGATE) for an accurate global representation, which may further contribute to its superior classification performance to SLAT and STAGATE^[11,13]^.

Synthetic data 2 presented a more complex scenario involving three classes of slices: class 1 emulates healthy tissue containing two cell types, class 2 includes an extra spatially separate “tumor” cell type, and class 3 represents tumor tissue with infiltration. When compared with other methods, PASSAGE also achieved the highest classification accuracy (**Figure S3**), with neat clustering of slice-level PASSAGE embeddings (**Figure S4, S5**), again demonstrating the effectiveness of PASSAGE in distinguishing highly heterogeneous spatial transcriptomic slices.

Importantly, the attention pooling layer in PASSAGE facilitates the straightforward identification of specific spatial regions within individual slices that contribute the most to phenotypic classification. As visualized by scaled attention scores in the attention pooling layer (**Methods**), PASSAGE precisely highlights spatial signatures designed into the simulation, in this case, cell type 2 which exhibits differential abundance and spatial distribution across phenotypic classes (**Figure 2E, S6**).

### PASSAGE effectively calls cancer-associated signatures within multiple heterogeneous datasets

To further demonstrate the performance of PASSAGE in real-world data, we compiled a large breast tumor dataset comprising 103 slices from 42 patients (**Table S1**) sourced from two experimental platforms: Spatial Transcriptomics (ST) and 10x Visium^[7,17–20]^. This dataset includes three phenotypic classes: 10 slices of normal breast tissue, 36 slices of conventional breast cancer tissue (positive for at least one of *ER, HER2, PR*), and 57 slices of triple-negative breast cancer tissue (**Figure 3A, Methods**). To ensure the generalizability of our model, we intentionally partitioned the training and testing sets by selecting slices from different experimental platforms and patients (**Methods**).

**Figure 3.**
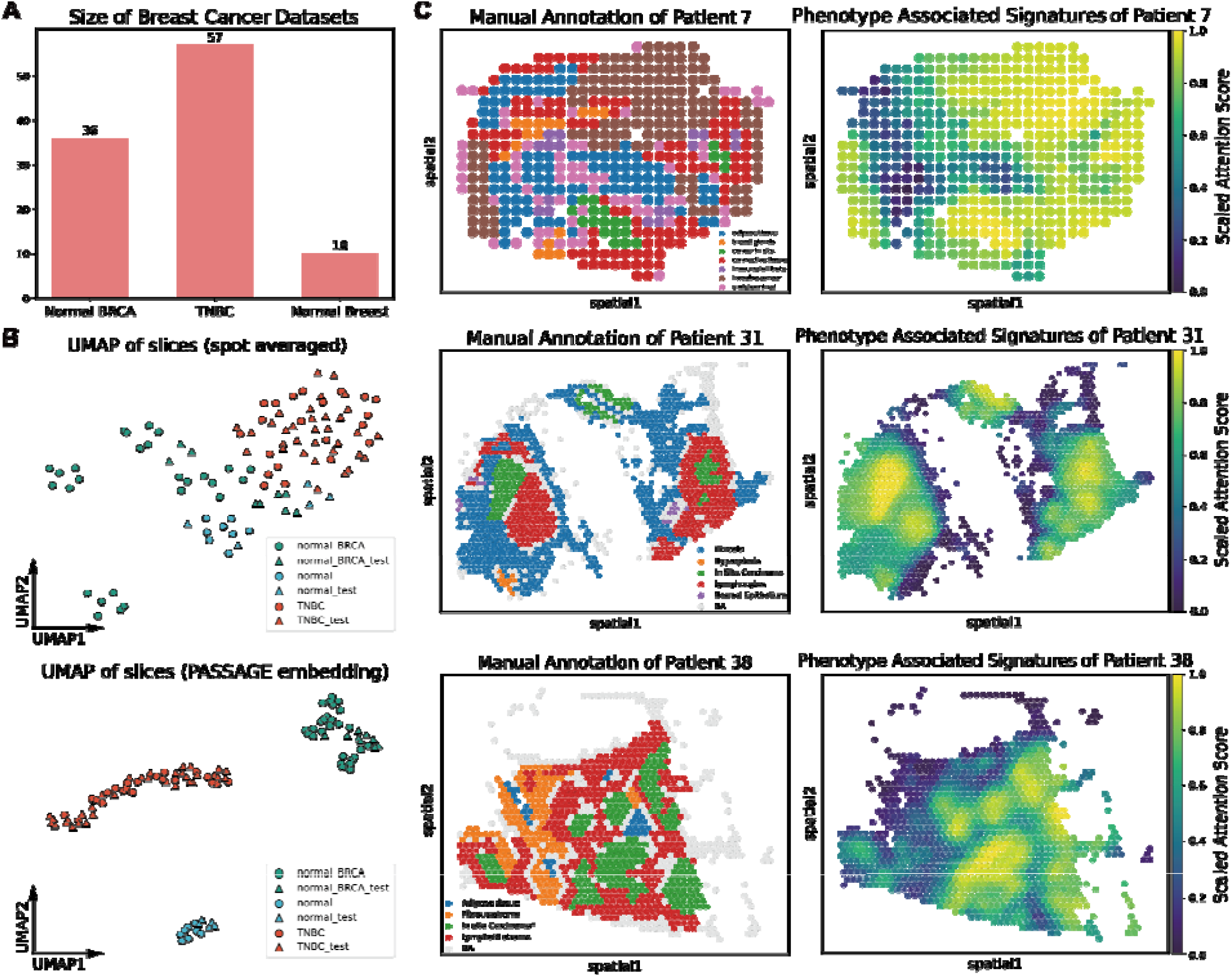
Evaluation of PASSAGE in the breast cancer dataset. A) Number of slices for each category in the breast cancer dataset. B) UMAP visualization of breast tissue slices obtained by simply averaging all spots (top) or using PASSAGE (bottom). C) Phenotype-associated spatial signatures identified by PASSAGE compared to manually annotated tissue regions for patient 7, 31 and 38.

PASSAGE consistently outperforms baseline methods in classifying the three phenotypic classes, achieving the highest accuracy (**Figure S7**). Notably, as previously mentioned, we specifically chose a test set consisting of breast tissue spatial slices from an experimental batch that had never been included in the training data. Due to batch effects, the three normal breast tissue slices from this batch are spatially closer to the triple-negative breast cancer slices from the same batch and relatively more distant from normal breast tissue slices collected from another experimental batch (**Figure S8**). However, PASSAGE effectively captured phenotype-associated representations during training, thereby accurately classifying the three normal breast tissue slices in the test set. Accordingly, PASSAGE also maintained accurate clustering performance (**Figure 3B, S9**), even when dealing with slices from different platforms, patients, and experimental batches that were never part of the training process, underscoring PASSAGE’s generalization capability and robustness in handling diverse and unseen data.

We then focused on the ability of PASSAGE to discover phenotype-related spatial signals on real data. In the breast tumor datasets, PASSAGE effectively identifies malignancy-associated spatial signatures despite their heterogeneous origins via its attention scores. For example, in patient-7 with conventional breast cancer (ER-positive, HER-2 positive), all invasive cancer tissue regions annotated manually by pathologists were accurately identified by PASSAGE, highlighting the strong correlation with the phenotype (**Figure 2E, S10**). Additionally, in patient-38 (triple-negative breast cancer), PASSAGE also successfully identified the dispersed tumor core regions in the slices (**Figure 3C**). Notably, in patient-31, also with triple-negative breast cancer, PASSAGE detected the disease-associated signatures, i.e., the tumor cells and a crucial tertiary lymphoid structure (TLS), although the Germinal Center (GC) of this TLS structure is not fully revealed in the HE-stained slices (**Figure S11, S12**). Furthermore, PASSGAE identified several lymphocyte-enriched foci on the right side of this spatial slice, pointing out a potential precursor area of incomplete TLS development.

To demonstrate the generality of PASSAGE, we further built a Squamous Cell Carcinoma (SCC) dataset, including 12 Oral Squamous Cell Carcinoma (OSCC) slices and 12 Cutaneous Squamous Cell Carcinoma (CSCC) slices^[21,22]^. We focused on malignancy-associated spatial signatures detected by PASSAGE in this dataset. **Figure 4A** illustrates the signature of CSCC patient-53 defined by PASSAGE. Compared to the manually annotated slices in the original study, PASSAGE accurately identified the entire tumor-associated regions.

**Figure 4.**
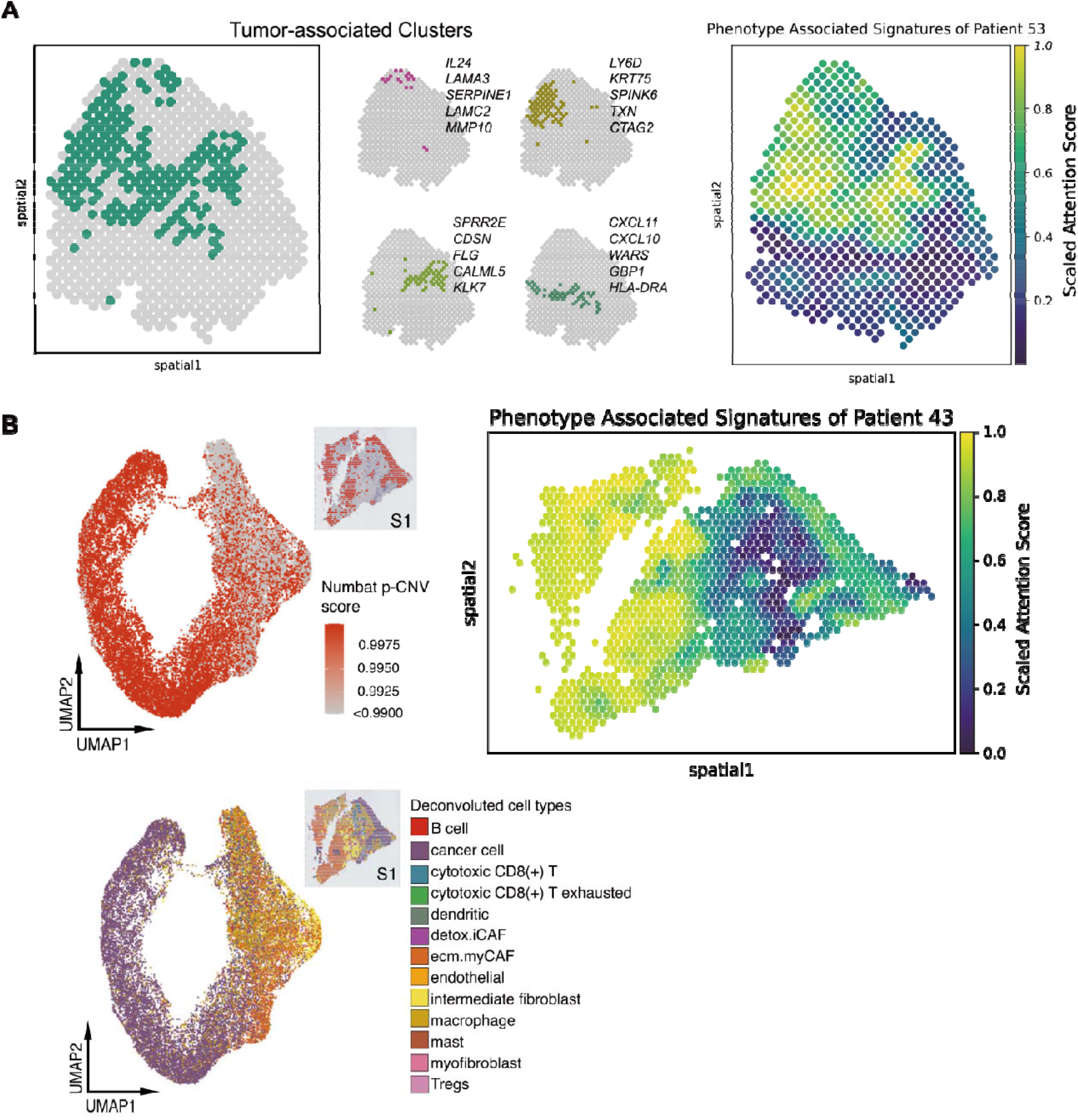
Phenotype-associated spatial signatures identified by PASSAGE in the SCC dataset. A) Phenotype-associated spatial signature identified by PASSAGE (right) compared to manually annotated tumor-associated clusters (left, from Figure S3 of Ji et.al.^[21]^) for patient 53 in the CSCC dataset. B) Numbat p-CNV score and deconvolution-based tumor-associated patterns from the original study (left, from Figure 1 of Arora et al.^[29]^) and PASSAGE-learned phenotype-associated spatial signatures for patients 43 in the OSCC dataset.

As for OSCC slices, compared to the p-CNV score inferred by Numbat and the deconvolution results from the original study, the phenotype-associated spatial signatures identified by PASSAGE in patient-43 from different patients significantly enriched in tumor-related regions (**Figure 4B**) ^[22,23]^. The rationality and biological interpretability of the phenotype-associated spatial signatures we identified can also be validated through the expression profiles of OSCC-specific marker genes (**Figure S13**).

These results indicate PASSAGE’s effectiveness in identifying biologically meaningful phenotype-associated spatial signatures, thereby facilitating researchers in discovering and deeply exploring the molecular features of pathological spatial slices with phenotype labels.

## Discussion

PASSAGE is designed as a global signature identification algorithm tuned for large-scale heterogeneous spatial slices. One essential challenge for PASSAGE is to learn a slice-wise representation for numerous within-slice spots, with biologically meaningful phenotype distinctions well preserved. Here, we introduced a dedicated attention pooling component for PASSAGE, to learn the contribution of individual cells/spots to the slice-level representation dynamically. A key advantage of the attention pooling component lies in its inherent capacity for adaptive information aggregation, which further enhances the interpretability of PASSAGE-identified signatures within the context of phenotype association.

Of note, the supervised nature of PASSAGE would effectively “encourage” the most parsimonious signature(s) for discriminating slices with distinct phenotypic groups. We noticed that, in real-world cases, such signatures could even be reduced to rather non-spatial ones, like the cell composition difference (e.g., the phenotype-associated signatures displayed in the benchmarking of synthetic data 2 are highly correlated with simulated cell type 2) as long as they can distinguish slices within different phenotypic groups effectively.

The rapid accumulation of spatial omics data enables a systematic reference atlas for comparative studies across developmental stages^[24]^, demographic populations^[25]^, as well as experimental perturbations^[26]^, analogous to what has been done for scRNA-seq^[27]^. We believe the PASSAGE, as an effective algorithm for calling phenotype-associated signatures globally, would be a valuable plus to the toolkits of both computational and experimental biologists. The whole package of PASSAGE, along with tutorials and demo cases, is available online at https://github.com/gao-lab/PASSAGE for the community.

## Methods

### Data preprocessing and spatial graph construction

We define a set of spatial slices as 𝒮 ={*S*^(*k*)^ ∣ *k =* 1,2,⋯, *N*} where *N* is the number of all training slices. Each slice is denoted as 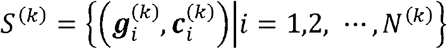, *N*^(*k*)^ is the number of spots in slice 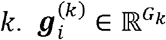 and 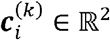 are the raw omics features (e.g., gene expression) and spatial coordinates of spot *i*,respectively.

For each spatial slice, we first removed low-quality spots with detected genes lower than 200. Then, raw expression values were normalized and log-transformed via scanpy (v1.9.6). Feature unification across slices is conducted in “outer” mode, which retains the union of detected genes across all spatial slices. Ultimately, spatial k-NN graphs for each slice are generated by torch-cluster (v1.6.3) and used as the input of PASSAGE, denoted as ℋ ={*h*^(*k*)^ ∣ *k =* 1,2,⋯, *N}*.

### Graph attention auto-encoder (GATE) module for learning spot-level embeddings

We adopt the GATE architecture in STAGATE as a spot-level embedding model, which in turn facilitates subsequent learning of slice-level embeddings. To reinforce the intrinsic information of each spot captured by the model, we introduced self-loops in the input spatial graph. For numerical stability, the output from the graph-encoder layer is normalized before being passed to the graph-decoder layer. The GATE module is trained by minimizing the mean squared error (MSE) reconstruction loss of the spot-level omics features. We denote the obtained spot-level embeddings as 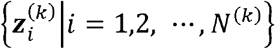. These GATE spot-level embeddings are trained and fixed before proceeding to train the slice-level embeddings (next section).

### Attention pooling for learning slice-level embeddings

To generate embeddings for entire slices, we propose the following attention pooling layer. The attention pooling layer boils down to a weighted sum of spot-level embeddings, where the weights are dynamically determined by an attention mechanism between all spots and an attention head^[28]^. To begin with, the spot-level embeddings 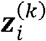 are first passed through a multi-level perceptron (MLP) to obtain 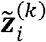 The attention head ***u***^(k)^ is computed by averaging the resulting embeddings across all spots in the slice, after applying a learnable linear transformation ***W***:

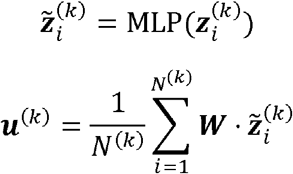

On top of that, we compute the attention weights based on the inner product ⟨ · ⟩ between each spot and the attention head above:

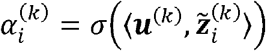

where is the sigmoid function. The attention score will be scaled to [ 0,1]interval to obtain the ‘scaled attention score’, which used for spatial signature visualization. Eventually, the slice-level embedding ***v***^(k)^ is calculated using the above attention weights:

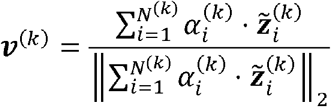

### Model optimization

Denote the categorical labels of slices *𝒸* ={*c*^(*k*)^ ∣ *k* = 1,2,⋯, *N*}. We randomly sampled triplet set *𝒱 =*{(*v*^(*i*)^, *v*^(*j*)^, *v*^(*k*)^) ❘ *c*^(i)^ = *c*^(*j*)^ ≠ *c*^(k)^,*i*,*j*,*k* = 1,2, ⋯, *N}* for training, where *v*^(*i*)^, *v*^(*j*)^ forms a positive pair and *v*^(*i*),^ *v*^(*k*)^ forms a negative pair. The size of triplet dataset ❘ 𝒱 ❘can be specified as a hyperparameter. Hence, the loss of multi-class contrastive learning is:

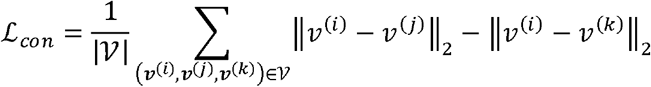

### Breast cancer and SCC dataset

For breast cancer dataset benchmarking, We collected 93 breast cancer slices and 10 corresponding healthy breast tissue slices from 4 studies^[17–20]^. We removed one of the spatial slices from patient-37 (denoted as ‘M10’ by Coutant et al.^[20]^), which was annotated as a TNBC sample with manual annotations from pathologists, due to the uncertainty regarding whether cells on the right hand side are genuine tumor cells or simply artifacts of tissue folding. For SCC dataset benchmarking, we collected 12 CSCC slices and 12 OSCC slices from 2 studies^[21,29]^. The data preprocessing process is consistent in all spatial slices as mentioned before.

### Synthetic datasets

Synthetic data 1 consisted of two classes of simulated spatial slices, each containing 160 slices. Each slice contained 2 cell types with different spatial distributions (**Figure 2A**). Spots of each cell type were generated according to an 8-dimensional multivariate normal distribution with mean µ_*i*_ ∼ Normal (3 ·**1**_8_, ***I***_8_),(*i* = 1,2) and variance 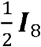. To increase the difficulty of classification, the proportions of the two cell types within each slice are the same.

Synthetic data 2 consisted of three classes of simulated spatial slices, each containing 25, 30, and 25 slices. Each slice contained 3 cell types with different spatial distributions (**Figure 2A**). Spots of each cell type were generated according to an 8-dimensional multivariate normal distribution with µ_*i*_ ∼ Normal (3 ·**1**_8_, ***I***_8_),(*i* = 1,2,3) and variance 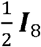.

### Benchmarking procedures

For both simulation and real-world datasets, we systematically quantified the performance against following baseline models: decision tree, k-NN, SVM, logistic regression, random forest, GAT classifier, STAGATE and SLAT, which were implemented using Python packages “sklearn” (v1.3.2), “torch_geometric” (v2.4.0), “STAGATE_pyG” and “scSLAT” (v0.2.1)^[11,13,30–32]^. All non-deep learning classification algorithms, we first averaged the omics features across all spots in each slice to obtain slice-level inputs, and the algorithms were run with their default settings. For STAGATE and SLAT, their learned unsupervised embeddings were first averaged across all spots in each slice, and then fed to a slice-level k-NN classifier. The GAT classifier constituted two graph convolutional layers followed by global average pooling across all spots in each slice and a fully connected layer for classification. The first and second GAT layers used single-head attention and the size of hidden layers were 64 and 16, respectively. For all runs of the PASSAGE model, we used the default hyperparameters (GATE hidden layer-1 size: 128, GATE hidden layer-2 size: 16, attention pooling layer size: 8 in simulation and 16 in real-world dataset, learning rate: 0.001, dropout probability: 0.3, AdamW weight decay: 5e-4, gradient norm clipping: 3). Both GAT and PASSAGE were trained for 10 epochs. All benchmarking methods were run with 10 different random seeds. All benchmarking tasks were accomplished on a server with Intel Xeon Platinum 8352V CPU and one NVIDIA RTX 4090 GPU.

## Data Availability

All real-world spatial transcriptomic datasets were downloaded from public databases, with accession numbers provided in **Table S1**^[17–22]^.

## Code Availability

The source code of PASSAGE is publicly available at https://github.com/gao-lab/PASSAGE.

## Supporting Information

Supporting Information is available from the Wiley Online Library or from the author.

## Acknowledgements

We thank Dr. G. Zhang for his helpful guidance on H&E staining slices scrutinization of breast tissue datasets, and J.W. Yao for her help in benchmark validation and code optimization. The research is supported by National Natural Science Foundation of China (grant no. 323B2017 to C.R.X, 32270854 to G.P.), the China Postdoctoral Science Foundation (grant no. 2023T160009 to Z.J.C.), the State Key Laboratory of Protein and Plant Gene Research and the Beijing Advanced Innovation Center for Genomics at Peking University, as well as the Changping Laboratory.

## Author contributions

G.G. and Z.J.C. conceived the study. G.G. Z.J.C and G.P. supervised the project. C.R.X. and C.K.G designed and actualized the computational framework, and conducted benchmarking and case studies with guidance by G.G., Z.J.C and G.P. C.K.G, C.R.X, G.P, Z.J.C and G.G. wrote this manuscript.

