## Supplementary Information for "Learning phenotype associated signature in spatial transcriptomics with PASSAGE"

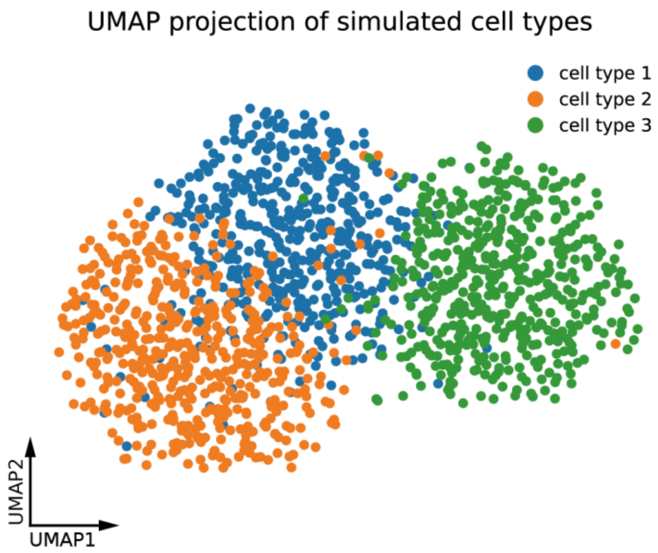


**Figure S1 Characteristics of simulated cell types.**

UMAP visualization of simulated spots in the spatial slices of synthetic data 1 & 2 (500 spots for each simulated cell type).

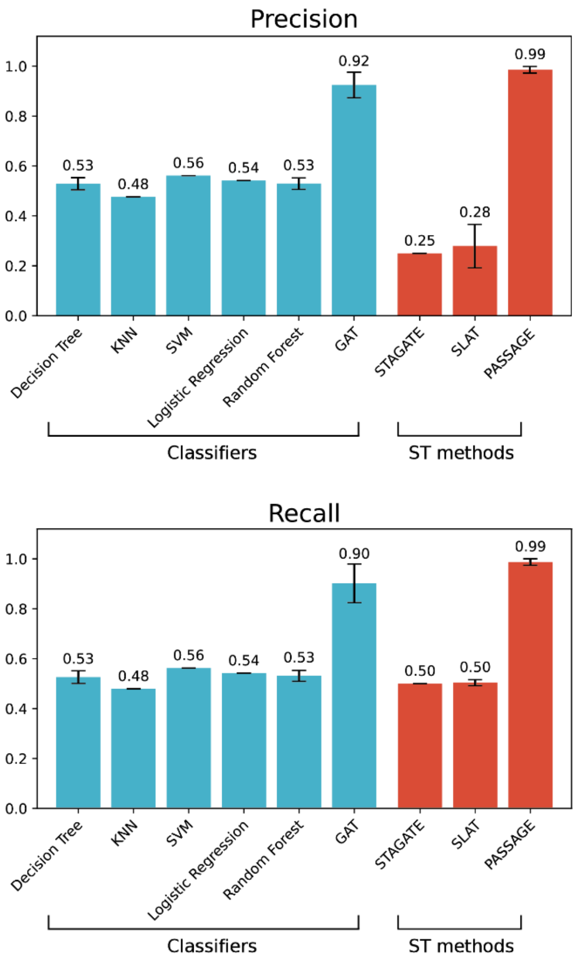


**Figure S2 Precision and recall of slice classification in synthetic data 1.**

The error bars indicate mean ± s.d.

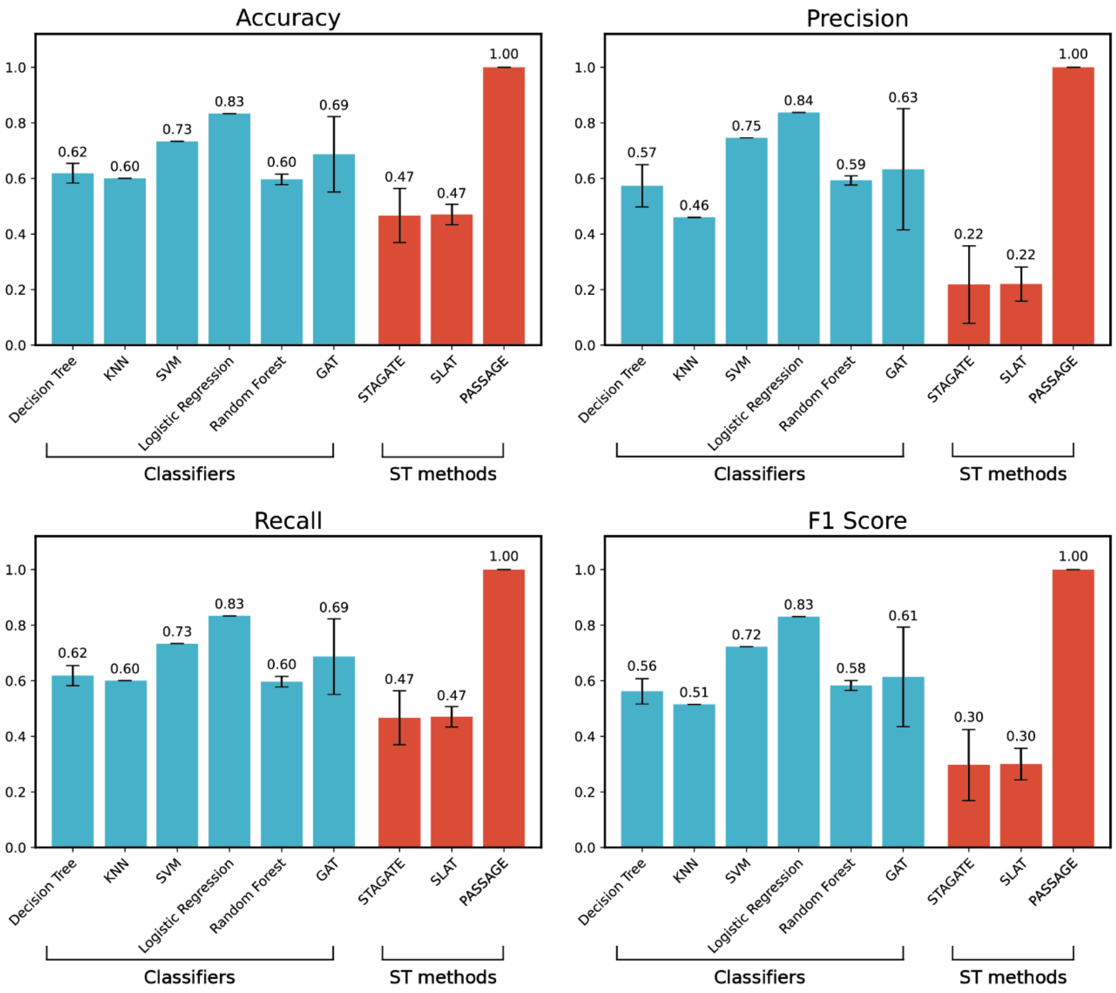


**Figure S3 Classification benchmarking in synthetic data 2.**

The error bars indicate mean ± s.d.

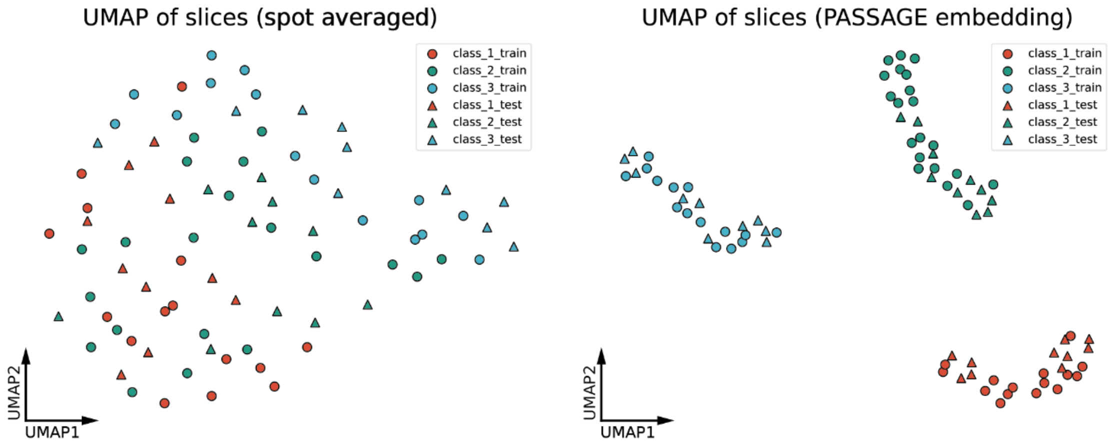


**Figure S4 UMAP visualization of slice-level embeddings in synthetic data 2.**

A) Slice embeddings obtained by simply averaging all spots. B) Slice embeddings obtained with PASSAGE.

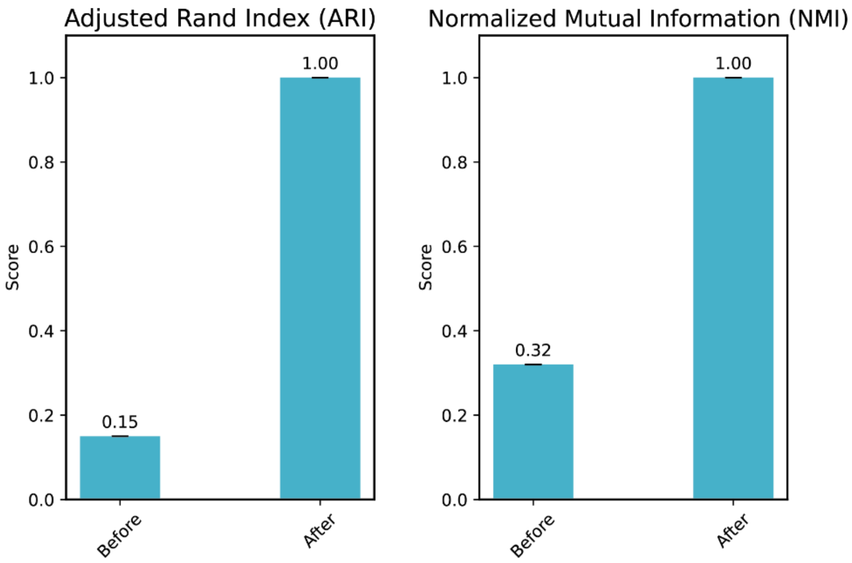


**Figure S5 Clustering metrics for the averaging-based embeddings and PASSAGE embeddings in synthetic data 2.**

The error bars indicate mean ± s.d.

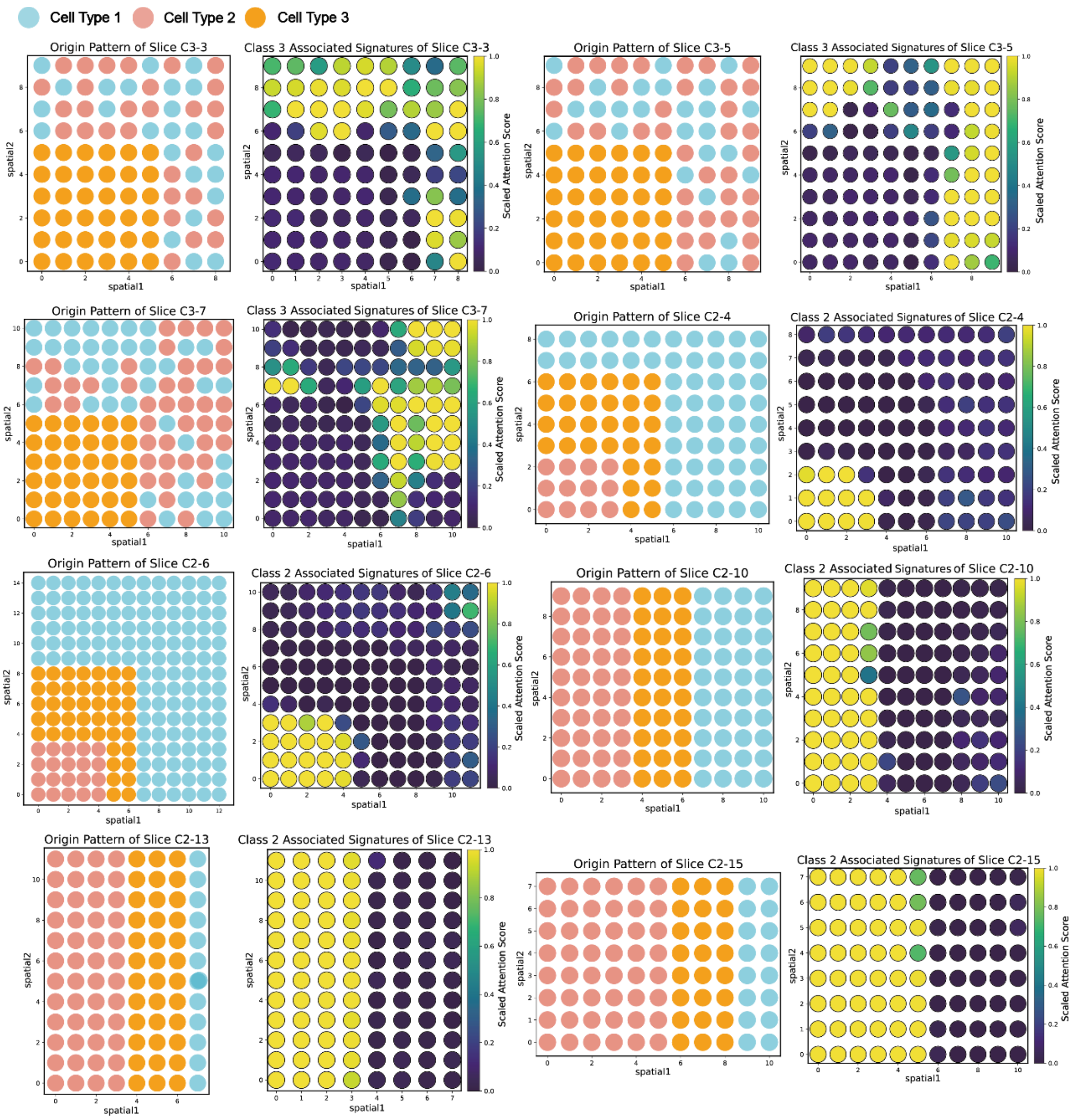


**Figure S6 Comparison of PASSAGE-identified phenotype-associated spatial signatures with ground-truth cell type annotations in synthetic data 2.**

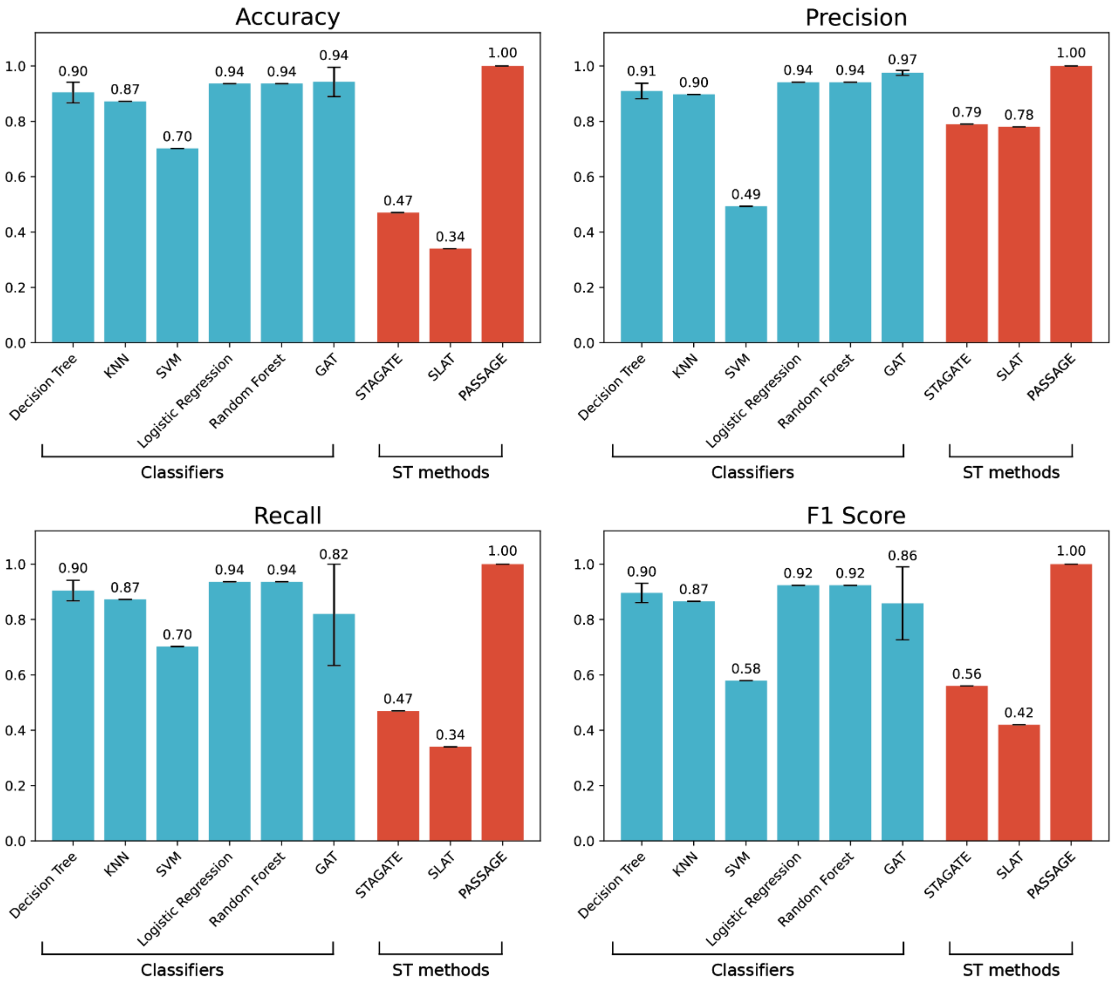


**Figure S7 Classification performance benchmarking in the breast cancer dataset.**

The error bars indicate mean ± s.d.

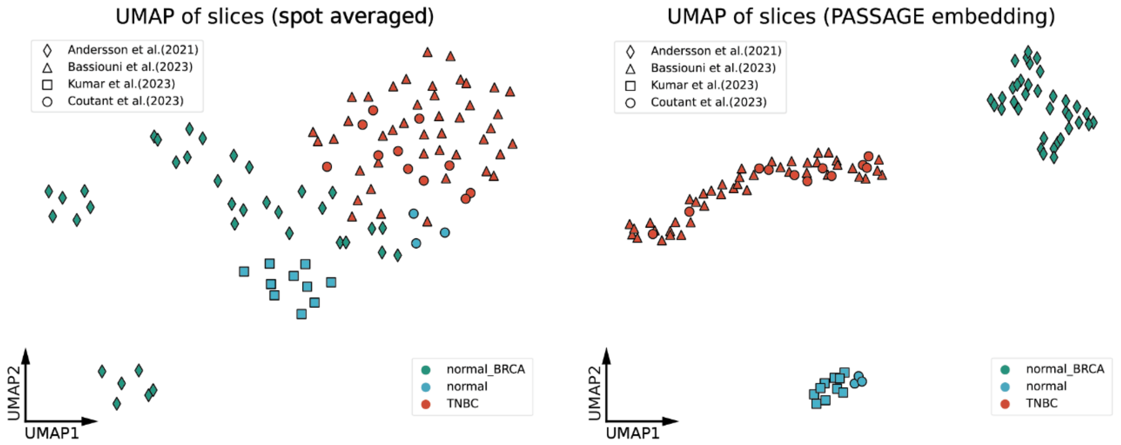


**Figure S8 UMAP visualization of slice embeddings in BRCA slices.**

Visualization of the original and PASSAGE-learned slice-level embedding distribution of BRCA slices labeled with dataset sources.

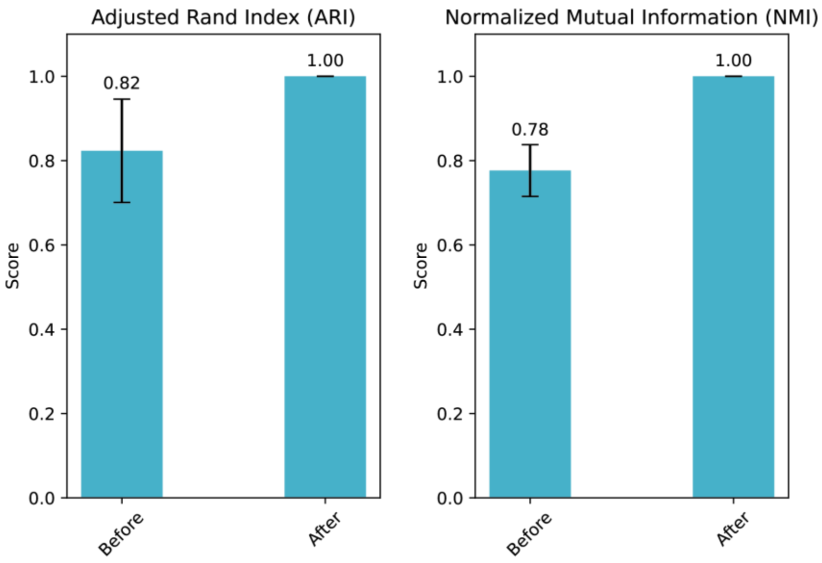


**Figure S9 Clustering metrics for the averaging-based embeddings and PASSAGE embeddings in the breast cancer dataset.**

The error bars indicate mean ± s.d.

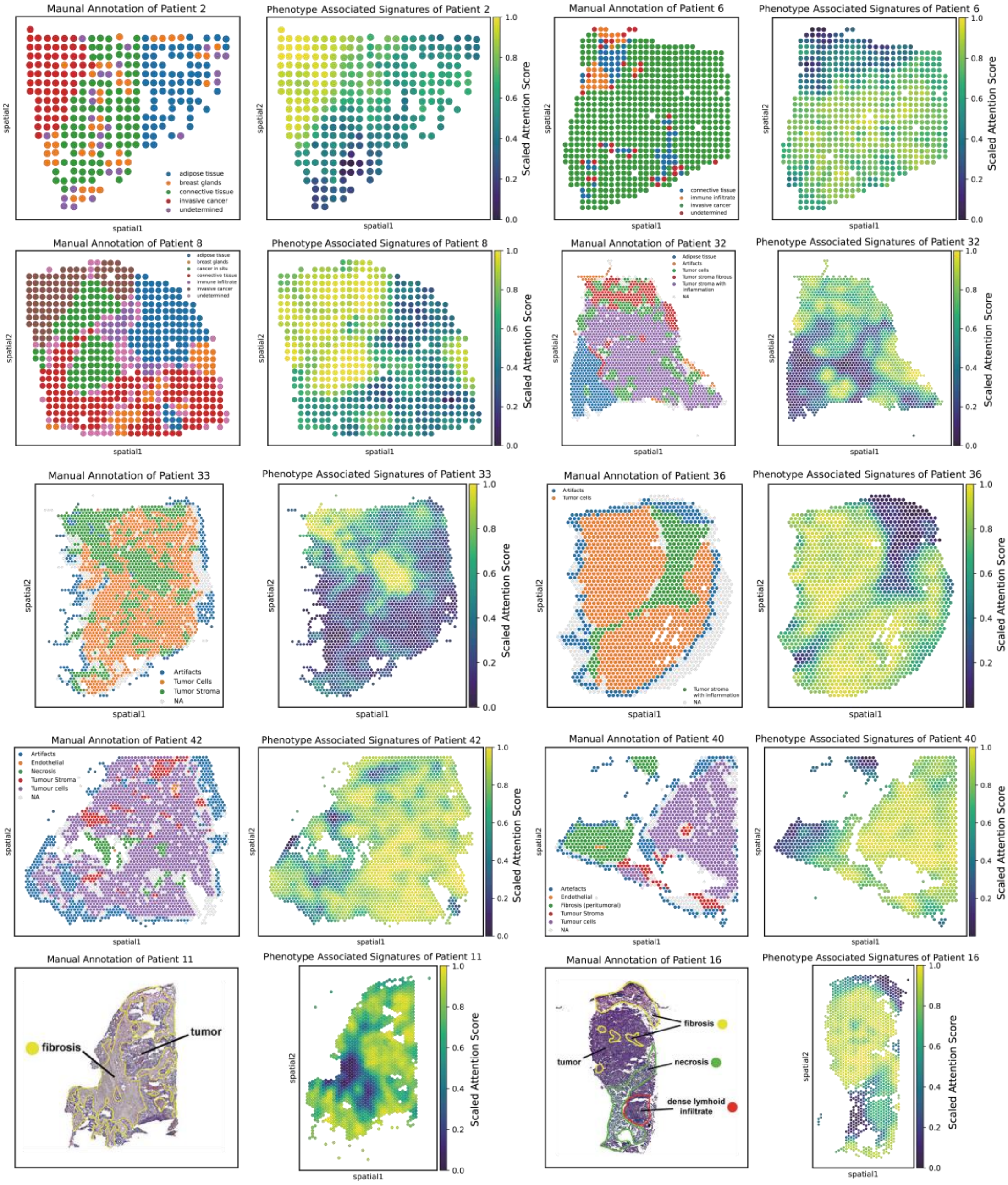


**Figure S10 Comparison of PASSAGE-identified phenotype-associated spatial signatures with manual annotations in the testing split of the breast cancer dataset.**

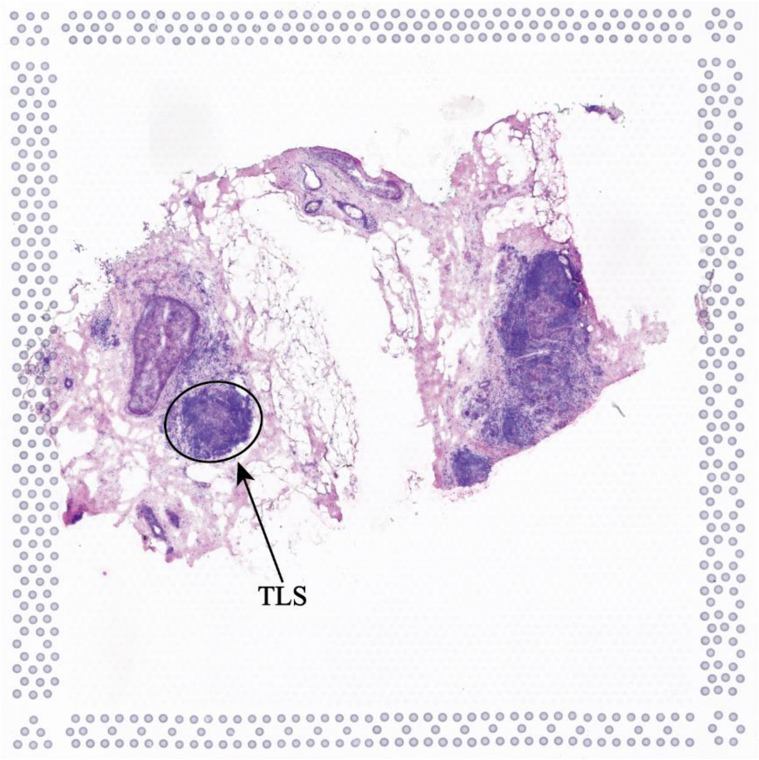


**Figure S11 H&E staining highlighting pre-mature TLS structure in patient 38.**

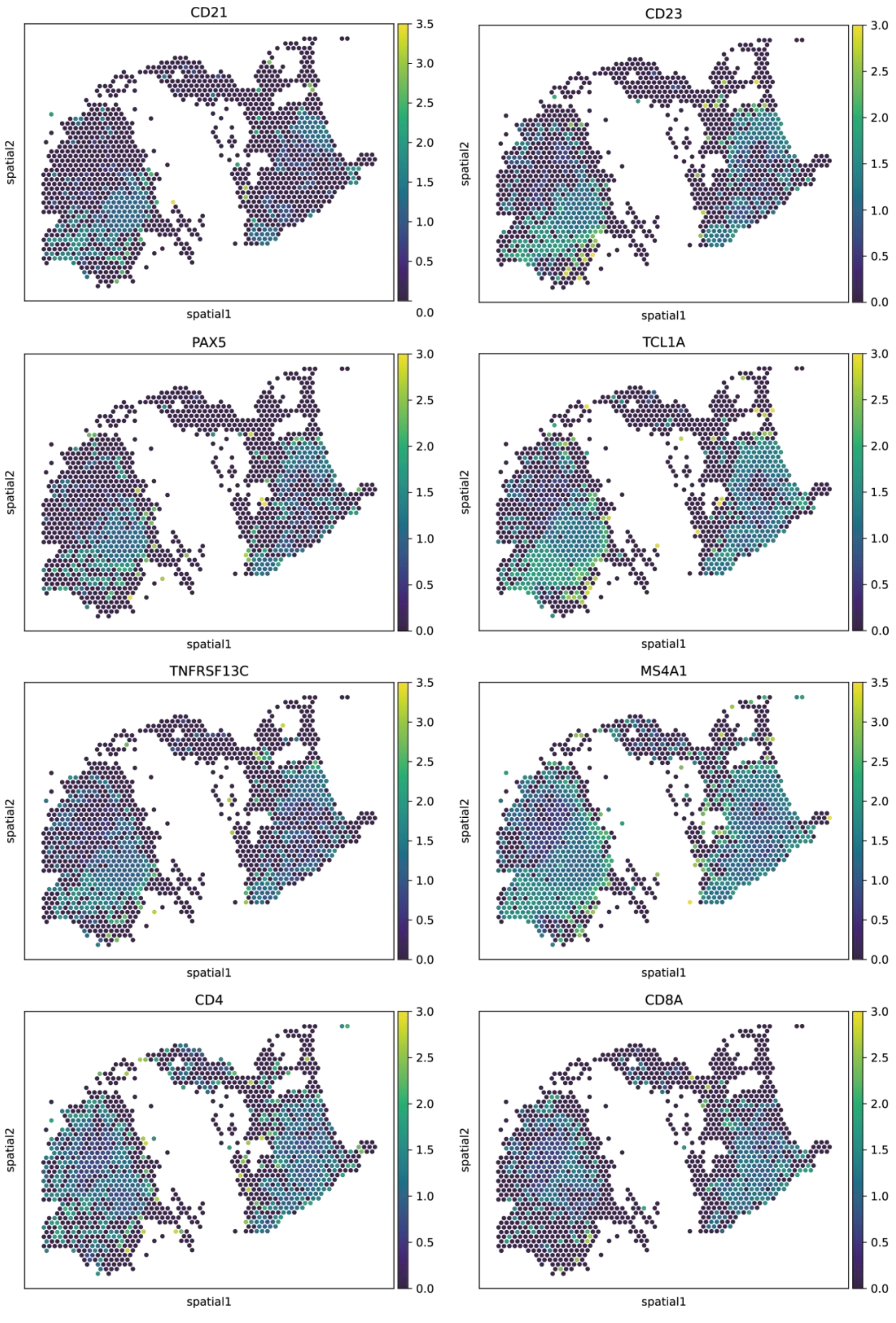


**Figure S12 Visualization of TLS-related markers in spatial slice of patient 38.** Visualization of various TLS-related markers in the spatial slice of patient 38. *CD21* and *CD23* are the cell type markers of follicular dendritic cells. *PAX5*, *TCL1A* and *TNFRSF13C* are the three gene signature of TLS. *MS4A1* is the marker of B cells. *CD4*, *CD8A* are the markers of T cells[1–3].

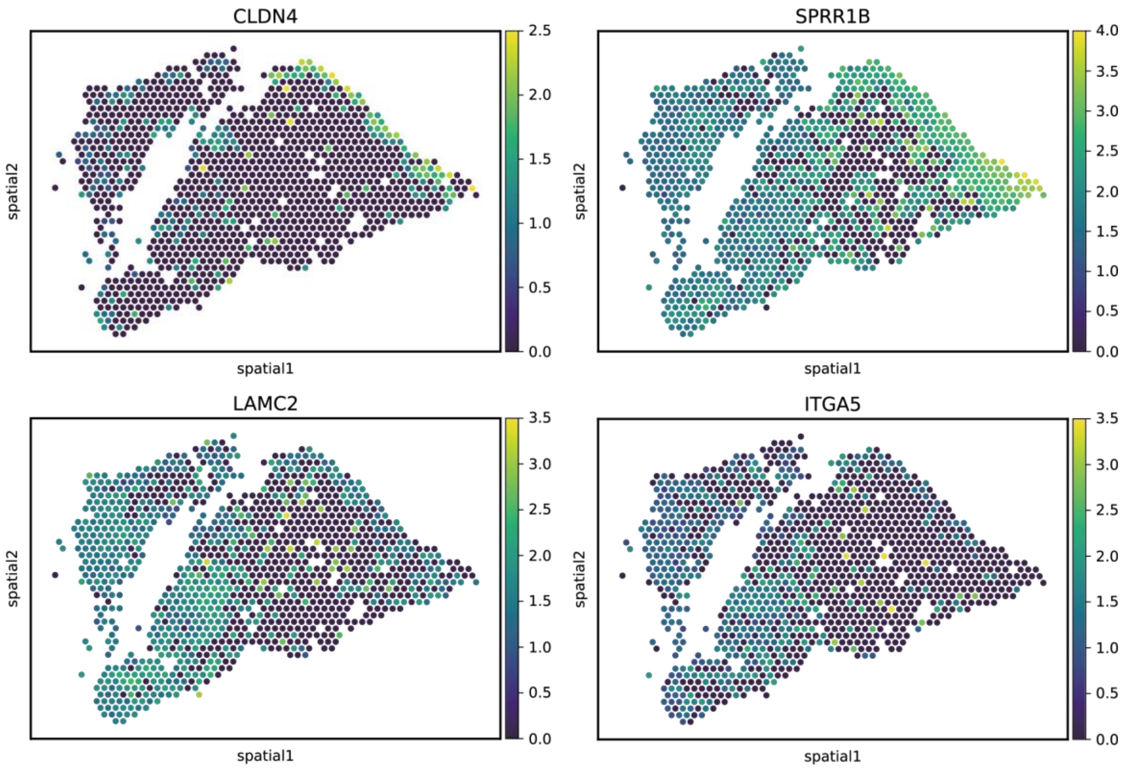


**Figure S13 Visualization of OSCC-related markers in spatial slice of patient 43.** Visualization of various OSCC-related markers in the spatial slice of patient 43. *CLDN4* and *SPRR1B* are tumor core markers previously validated in head and neck cancer[4]. *LAMC2* and *ITGA5* are tumor-associated DEGs from Arora et al[5].

| Patient ID | Phenotype annotation | Number of slices | Technology | Data sources | Reference |
|---|---|---|---|---|---|
| Patient 1 | ER negative, PR negative, HER2 positive, Invasive Breast Carcinoma | 6 | ST | https://github.com/almaan/her2st/ | [6] |
| Patient 2 | ER negative, PR positive, HER2 positive, Invasive Breast Carcinoma | 6 | ST | https://github.com/almaan/her2st/ | [6] |
| Patient 3 | ER negative, PR negative, HER2 positive, Invasive Breast Carcinoma | 6 | ST | https://github.com/almaan/her2st/ | [6] |
| Patient 4 | ER negative, PR negative, HER2 positive, Invasive Breast Carcinoma | 6 | ST | https://github.com/almaan/her2st/ | [6] |
| Patient 5 | ER negative, PR negative, HER2 positive, Invasive Breast Carcinoma | 3 | ST | https://github.com/almaan/her2st/ | [6] |
| Patient 6 | ER negative, PR negative, HER2 positive, Invasive Breast Carcinoma | 3 | ST | https://github.com/almaan/her2st/ | [6] |
| Patient 7 | ER negative, PR negative, HER3 positive, Invasive Breast Carcinoma | 3 | ST | https://github.com/almaan/her2st/ | [6] |
| Patient 8 | ER negative, PR negative, HER4 positive, Invasive Breast Carcinoma | 3 | ST | https://github.com/almaan/her2st/ | [6] |
| Patient 9 | Triple Negative Breast Cancer | 2 | 10x Visium | GSE210616 | [7] |
| Patient 10 | Triple Negative Breast Cancer | 4 | 10x Visium | GSE210616 | [7] |
| Patient 11 | Triple Negative Breast Cancer | 4 | 10x Visium | GSE210616 | [7] |
| Patient 12 | Triple Negative Breast Cancer | 2 | 10x Visium | GSE210616 | [7] |
| Patient 13 | Triple Negative Breast Cancer | 4 | 10x Visium | GSE210616 | [7] |
| Patient 14 | Triple Negative Breast Cancer | 4 | 10x Visium | GSE210616 | [7] |

| Patient ID | Phenotype annotation | Number of slices | Technology | Data sources | Reference |
|---|---|---|---|---|---|
| Patient 15 | Triple Negative Breast Cancer | 4 | 10x Visium | GSE210616 | [7] |
| Patient 16 | Triple Negative Breast Cancer | 4 | 10x Visium | GSE210616 | [7] |
| Patient 17 | Triple Negative Breast Cancer | 4 | 10x Visium | GSE210616 | [7] |
| Patient 18 | Triple Negative Breast Cancer | 3 | 10x Visium | GSE210616 | [7] |
| Patient 19 | Triple Negative Breast Cancer | 4 | 10x Visium | GSE210616 | [7] |
| Patient 20 | Triple Negative Breast Cancer | 4 | 10x Visium | GSE210616 | [7] |
| Patient 21 | Paracancerous Tissue (normal section) | 1 | 10x Visium | GSE195665 | [8] |
| Patient 22 | Paracancerous Tissue (normal section) | 1 | 10x Visium | GSE195665 | [8] |
| Patient 23 | Normal Breast Tissue | 1 | 10x Visium | GSE195665 | [8] |
| Patient 24 | Normal Breast Tissue | 1 | 10x Visium | GSE195665 | [8] |
| Patient 25 | Normal Breast Tissue | 1 | 10x Visium | GSE195665 | [8] |
| Patient 26 | Normal Breast Tissue | 1 | 10x Visium | GSE195665 | [8] |
| Patient 27 | Normal Breast Tissue | 1 | 10x Visium | GSE195665 | [8] |
| Patient 28 | Normal Breast Tissue | 1 | 10x Visium | GSE195665 | [8] |
| Patient 29 | Normal Breast Tissue | 1 | 10x Visium | GSE195665 | [8] |

| Patient ID | Phenotype annotation | Number of slices | Technology | Data sources | Reference |
|---|---|---|---|---|---|
| Patient 30 | Normal Breast Tissue | 1 | 10x Visium | GSE195665 | [8] |
| Patient 31 | Paracancerous Tissue (normal section) | 3 | 10x Visium | GSE213688 | [9] |
| Patient 32 | Triple Negative Breast Cancer | 1 | 10x Visium | GSE213688 | [9] |
| Patient 33 | Triple Negative Breast Cancer | 1 | 10x Visium | GSE213688 | [9] |
| Patient 34 | Triple Negative Breast Cancer | 1 | 10x Visium | GSE213688 | [9] |
| Patient 35 | Triple Negative Breast Cancer | 1 | 10x Visium | GSE213688 | [9] |
| Patient 36 | Triple Negative Breast Cancer | 1 | 10x Visium | GSE213688 | [9] |
| Patient 37 | Paracancerous Tissue (normal section) | 1 | 10x Visium | GSE213688 | [9] |
| Patient 38 | Triple Negative Breast Cancer | 1 | 10x Visium | GSE213688 | [9] |
| Patient 39 | Triple Negative Breast Cancer | 1 | 10x Visium | GSE213688 | [9] |
| Patient 40 | Triple Negative Breast Cancer | 1 | 10x Visium | GSE213688 | [9] |
| Patient 41 | Triple Negative Breast Cancer | 1 | 10x Visium | GSE213688 | [9] |
| Patient 42 | Triple Negative Breast Cancer | 1 | 10x Visium | GSE213688 | [9] |
| Patient 43 | Oral Squamous Cell Carcinoma | 1 | 10x Visium | GSE208253 | [5] |
| Patient 44 | Oral Squamous Cell Carcinoma | 1 | 10x Visium | GSE208253 | [5] |

| Patient ID | Phenotype annotation | Number of slices | Technology | Data sources | Reference |
|---|---|---|---|---|---|
| Patient 45 | Oral Squamous Cell Carcinoma | 1 | 10x Visium | GSE208253 | [5] |
| Patient 46 | Oral Squamous Cell Carcinoma | 1 | 10x Visium | GSE208253 | [5] |
| Patient 47 | Oral Squamous Cell Carcinoma | 1 | 10x Visium | GSE208253 | [5] |
| Patient 48 | Oral Squamous Cell Carcinoma | 1 | 10x Visium | GSE208253 | [5] |
| Patient 49 | Oral Squamous Cell Carcinoma | 2 | 10x Visium | GSE208253 | [5] |
| Patient 50 | Oral Squamous Cell Carcinoma | 2 | 10x Visium | GSE208253 | [5] |
| Patient 51 | Oral Squamous Cell Carcinoma | 1 | 10x Visium | GSE208253 | [5] |
| Patient 52 | Oral Squamous Cell Carcinoma | 1 | 10x Visium | GSE208253 | [5] |
| Patient 53 | Oral Squamous Cell Carcinoma | 3 | 10x Visium | GSE208253 | [5] |
| Patient 54 | Cutaneous Squamous Cell Carcinoma | 3 | 10x Visium | GSE144240 | [10] |
| Patient 55 | Cutaneous Squamous Cell Carcinoma | 3 | 10x Visium | GSE144240 | [10] |
| Patient 56 | Cutaneous Squamous Cell Carcinoma | 3 | 10x Visium | GSE144240 | [10] |

**Table S1 Collected Spatial Transcriptomics Datasets.**

Spatial transcriptomics datasets utilized for training or testing our model, consisting of 104 slices from 56 patients. Datasets are publicly available via listed URLs or GEO accession number.
